# Interoception Network in the Rat Brain

**DOI:** 10.64898/2026.02.08.704721

**Authors:** Fatimah M. Alkaabi, Xiaokai Wang, Zhongming Liu

**Affiliations:** Department of Biomedical Engineering, University of Michigan, Ann Arbor, MI, USA; Department of Electrical Engineering and Computer Science, University of Michigan, Ann Arbor, MI, USA; Department of Biomedical Engineering, Imam Abdulrahman bin Faisal University, Dammam, Saudi Arabia

**Keywords:** Interoception, Resting State fMRI, Brain-Body Interaction, Functional Connectivity

## Abstract

The brain is never truly at rest. Even in the absence of external stimuli, the central nervous system continuously monitors and regulates visceral organs. However, it remains unclear to what extent this interoceptive signaling shapes the brain’s resting state networks. Here, we tested the hypothesis that resting state functional connectivity is actively sustained by visceral inputs. Using functional magnetic resonance imaging in 34 anesthetized rats, we identified a cohesive “interoception network” characterized by dense, reciprocal functional connectivity among the dorsal vagal complex, thalamus, hypothalamus, amygdala, insula, and cingulate cortex. The topology of this network was not static but dynamically coupled to the physiological state. Functional integration was robust in the digestive (fed) phase but significantly attenuated in the inter-digestive (fasted) phase. The network relied on the integrity of the vagus nerve. When vagal signaling was eliminated by bilateral cervical vagotomy, the network effectively disintegrated. These findings demonstrate that resting state functional connectivity is not an intrinsic property of the isolated brain but an embodied phenomenon that relies on continuous neural signaling between the brain and viscera.

## Introduction

Given that perception, cognition, and behavior are the most observable aspects of human life, neuroscience has historically focused on studying how the brain interacts with the environment. Typical experimental paradigms use external stimuli or cognitive tasks to interrogate brain function, viewing the brain as either a “reactive system” or a “thinking machine” in isolation from the body. However, this predominant focus on “exteroception” neglects a more fundamental biological imperative: the brain must continuously sense and regulate the body’s internal organs and systems through “interoception” (Craig, 2002, 2003; Chen et al., 2021). From an evolutionary perspective, the ancient (subcortical) brain is largely dedicated to interoception to ensure survival (Saper, 2002); as the neocortex emerges, it further extends interoceptive processes to contextualize perception, behavior, emotion, and cognition (Critchley & Harrison, 2013; Barrett & Simmons, 2015; Critchley & Garfinkel, 2017; Azzalini et al., 2019; Engelen et al., 2023).

Interoception is particularly relevant to the so-called “resting state” (Raichle, 2011). When interactions with the external environment are minimized, the brain remains actively engaged with visceral organs (Liu et al., 2023). Spontaneous brain activity, observable with functional magnetic resonance imaging (fMRI), exhibits robust spatial patterns of temporal correlations (Biswal et al., 1995; Fox & Raichle, 2007). These correlations are widely referred to as “functional connectivity” that delineates the brain’s intrinsic networks (Greicius et al., 2003; Yeo et al., 2011; Smith et al., 2013). These networks are often interpreted through a cognitive lens (Laird et al., 2011), such as mind wandering (Mason et al., 2007) or memory consolidation (Tambini et al., 2010). However, such interpretations are incomplete, given that similar network patterns persist across species (Lu et al., 2012) and even in unconscious states, such as sleep (Horovitz et al., 2008) or under anesthesia (Vincent et al., 2007). This leads us to hypothesize that resting state functional connectivity likely reflects a more primitive physiological origin and is actively shaped by interoceptive signaling between the brain and viscera (Liu et al., 2023). However, interoceptive influence on resting state functional connectivity has rarely been investigated. Physiological fluctuations, especially those of cardiac and respiratory activity, are often viewed as nuisance sources of vascular or mechanical “noise” to the fMRI signal (Birn et al., 2006; Shmueli et al., 2007). Common preprocessing pipelines remove any variance associated with physiological signals (Glover et al., 2000; Birn et al., 2008) (Chang et al., 2009; Kasper et al., 2017). This practice, however, risks eliminating neural signals that reflect the interoceptive processing of bodily states (Chen et al., 2020; Liu, 2016; Bright et al., 2017; Liu et al., 2023).

Indeed, anatomical studies have uncovered the peripheral and central pathways that support interoception (Craig, 2002; Saper, 2002; Rinaman, 2010; Browning & Travagli, 2014; Gehrlach et al., 2020; Gasparini et al., 2020; Berntson & Khalsa, 2021). In particular, the vagus nerve transmits sensory information from visceral organs to the brain via afferent fibers and conveys motor commands from the brain to visceral organs via efferent fibers (Berthoud & Neuhuber, 2000; Prescott & Liberles, 2022; Bassi et al., 2022; Powley, 2021). These sensory and motor signals converge at the dorsal vagal complex (DVC) in the lower brainstem to exert autonomic, reflexive control over organ physiology (Travagli & Anselmi, 2016). From the DVC, sensory information ascends to the parabrachial nucleus, periaqueductal gray, hypothalamus, and amygdala, and is subsequently relayed via the thalamus to the insular cortex (Saper, 2002; Rinaman, 2010). This structural pathway enables interoceptive processing through dynamic neuronal interactions among its connected regions. However, whether and to what extent interoceptive processing accounts for resting state functional connectivity remains unknown.

Our study aimed to address this question by systematically assessing the interoceptive influence on resting state functional connectivity in anesthetized rats. We focused on a set of core regions situated along the central interoceptive pathway as established by prior studies (Saper, 2002; Rinaman, 2010; Browning & Travagli, 2014; Gehrlach et al., 2020; Gasparini et al., 2020; Berntson & Khalsa, 2021). Using resting state fMRI, we measured the functional connectivity among these regions and tested whether they formed a cohesive network driven by interoceptive signaling. Our hypothesis was that if this network is attributable to interoception, its functional connectivity must be sensitive to changes in interoceptive signals that visceral organs send to the brain via peripheral nerves. To test this hypothesis, we used two experimental manipulations to modulate interoceptive signals either at the visceral source or along the peripheral pathway. First, we manipulated the gastrointestinal (GI) tract by separating its physiological state into the digestive (fed) versus inter-digestive (fasted) phase. These phases are known to engage highly distinct patterns of GI physiology and gut-brain interaction: the digestive phase is characterized by postprandial distension, GI motility, nutrient handling and absorption (Goyal et al., 2019; X. Wang et al., 2024), whereas the inter-digestive phase is characterized by long periods of motor quiescence and intermittent contractions (also known as the migrating motor complex (Deloose et al., 2012)). This manipulation specifically targeted the GI tract, while avoiding cardiac and respiratory confounds that could complicate the interpretation of the fMRI signal (Cao et al., 2022). Second, we performed bilateral cervical vagotomy to surgically eliminate vagal signaling between the brain and thoracic and abdominal viscera (Cao et al., 2022). This surgical manipulation disconnected both vagal afferents and efferents, compromising both organ-to-brain sensory transmission and brain-to-organ parasympathetic control. Together, these manipulations allowed us to examine whether resting state functional connectivity is a self-sustained central phenomenon or whether it relies on brain-body interaction through neuronal signaling via peripheral neural pathways.

## Materials and Methods

### Animals and Experimental Design

Experiments were performed on a total of 34 adult rats (average weight: 300 ± 50 g; 25 male Sprague-Dawley (SD) and 9 female Fischer). All animals were housed in a temperature-controlled environment (21 ± 1*^◦^*C) with a 12-hour light/dark cycle (lights on from 6 AM to 6 PM). Animal care and experimental procedures were approved by the Institutional Animal Care and Use Committee (IACUC) and the Laboratory Animal Program at Purdue University.

Animals were assigned to three experimental cohorts to identify the interoception network (“baseline” cohort), test its dependence on the feeding state (“state-dependence” cohort), and test its dependence on vagal integrity (“vagotomy” cohort). The baseline cohort (*n* = 22; 9 female Fisher and 13 male SD) maintained *ad libitum* access to food and water. Data from this cohort were used to map the baseline resting state functional connectivity, identify the interoception network, and characterize its functional topology. The state-dependence cohort (*n* = 7 male SD) underwent dietary habituation such that animals could voluntarily consume a controlled test meal after an overnight fast. This cohort allowed us to perform fMRI experiments under controlled fed versus fasted conditions. The vagotomy cohort (*n* = 5 male SD) underwent both the dietary habituation and bilateral cervical vagotomy. This cohort allowed us to perform fMRI experiments under the fed condition in the absence of vagal signaling.

These datasets were originally collected in our previous studies (Liu & Cao, 2020; Cao et al., 2022) and were re-analyzed for the new objective of this study. In the baseline cohort, data from *n* = 9 female Fisher rats were from a carefully collected and curated dataset established for our exploratory discovery and inquiry of resting state networks (Liu & Cao, 2020), whereas data from *n* = 13 male SD rats were used for replication. Since there were no significant differences in resting state networks between Fisher and SD rats in the baseline cohort, data were pooled in the results reported herein. Note that from only male SD rats were the data used to test how resting state networks depended on the vagal integrity and feeding state. The current study was not designed or powered to test the effects of sex or strain, which were conflated in this study.

### Dietary Habituation

Using an established protocol for dietary habituation (Lu et al., 2017), we trained animals to rapidly and voluntarily consume a standardized test meal immediately following an overnight fast, providing naturalistic and controlled conditions in terms of gastrointestinal physiology (X. Wang et al., 2026) and gut-brain interaction (Cao et al., 2022). The test meal consisted of a fixed quantity (5 g) of palatable gel diet (DietGel Recovery, ClearH2O, ME, USA). The habituation period lasted for 7 days. During the first two days (acclimation), animals were provided with a small amount of the gel diet in their home cages alongside standard chow. During days 3–7 (scheduled feeding), animals were switched to a restricted feeding schedule; they were fasted overnight (18 hours) and then presented with 5 g of the gel diet. This repeated exposure preconditioned the animals to consume the entire portion rapidly upon presentation.

### Fasting and Feeding

On the day of the experiment, all animals were fasted for 18 hours. For the “fed” condition, animals were presented with 5 g of the test meal and allowed to consume it voluntarily. Animals completed the meal in about 5 minutes; otherwise the meal was taken away. Anesthesia induction began immediately after the meal. For the “fasted” condition, animals remained food-deprived and proceeded directly to anesthesia induction without access to the test meal. This design ensured that the sole variable differing between these two conditions was the meal intake, while controlling for potential confounds such as stress, circadian rhythm, and metabolic history.

### Cervical Vagotomy

For animals in the vagotomy cohort (*n* = 5), a bilateral cervical vagotomy was performed immediately following the test meal and prior to MRI. Anesthesia was induced with 5% isoflurane and maintained at 1.5–2.5% in oxygen-enriched air. The animal was positioned supine, and a midline incision (approximately 1.5 cm) was made in the anterior neck. The cervical vagus nerve was identified, isolated from the common carotid artery, and transected bilaterally (left side first). Following the vagotomy, the incision was closed with sterile sutures, and then the animal was positioned for fMRI acquisition. The vagotomy was completed about 15 minutes after the meal.

Since the left vagus was consistently transected immediately prior to the right vagus, there was a brief interval for a few minutes that the right vagus nerve was intact while the left vagus nerve was transected. During this brief interval, transient asymmetric signaling could theoretically occur through the right vagus nerve. However, it is unlikely that any right-lateralized neural events during this brief surgical window would have influenced subsequent imaging sessions that started at least half an hour later during a postprandial window that lasted over one hour.

### Anesthesia

All animals were scanned under comparable physiological and anesthetic conditions. Anesthesia was initially induced with 5% isoflurane in oxygen (1 L/min) and reduced to 1.5–2.5% during animal preparation. Once positioned in the scanner, a bolus of dexmedetomidine (0.0125 mg/kg) was administered subcutaneously at 1 L/min. About 15 minutes after the bolus, a continuous subcutaneous infusion of dexmedetomidine was initiated at a rate of 0.0125 mg/kg/h, while isoflurane was lowered to 0.1–0.5%. To maintain a stable depth of anesthesia, the infusion rate was increased by 0.0125 mg/kg/h for every additional hour of scanning.

### Physiological Monitoring

Physiological parameters were continuously monitored using a pulse oximeter (MouseSTAT, Kent Scientific, Torrington, CT, USA) for heart rate and SpO_2_ and a physiology monitoring system (Model 1030, SA Instruments, Stony Brook, NY, USA) for respiration and temperature. Body temperature was maintained at 36.5–37.5*^◦^*C using a feedback-controlled warm air heating system. FMRI data acquisition started when physiological parameters stabilized within the target ranges: heart rate 250–350 beats/min, respiratory rate 20–60 breaths/min, and SpO_2_ *>* 96%.

### MRI Acquisition

MRI experiments were conducted on a 7-Tesla small-animal MRI system (BioSpec 70/30, Bruker, Billerica, MA, USA). The system was equipped with an 86-mm inner diameter volume transmit coil and a quadrature surface receive coil. Animals were positioned prone on a custom-designed holder, with the head secured using ear bars and a bite bar to minimize motion artifacts. Anatomical images were acquired using a 2-D Rapid Acquisition with Relaxation Enhancement (RARE) sequence with the following parameters: repetition time (TR) = 5804.6 ms, effective echo time (TE) = 32.5 ms, echo spacing = 10.83 ms, RARE factor = 8, flip angle (FA) = 90*^◦^*. The anatomical volume consisted of 50 slices with a voxel size of 0.125 × 0.125 × 0.5 mm^3^, covering the entire brain. Blood oxygenation level dependent (BOLD) fMRI data were subsequently acquired using a 2-D single-shot gradient-echo echo-planar imaging (GE-EPI) sequence. Parameters were set to: TR = 1000 ms, TE = 16.5 ms, FA = 55*^◦^*, matrix size = 64 × 64, field of view = 32 × 32 mm^2^, in-plane resolution = 0.5 × 0.5 mm^2^, and slice thickness = 1 mm (25 slices). Each functional run consisted of 1800 volumes, lasting 30 minutes. Animals in the fed and vagotomy cohorts underwent, two runs of fMRI (total time: 60 min), interleaved with dynamic MRI of the gastrointestinal (GI) tract, were performed simultaneously with body-surface electrogastrogram (EGG) recordings. Both EGG and GI MRI data were outside the scope of this paper. Animals in the fasted cohort underwent four consecutive fMRI runs (total time: 120 min) to capture the potentially higher variability in the fasted state.

### MRI Preprocessing

A customized preprocessing pipeline was applied to all datasets using MATLAB (The MathWorks, Inc., 2024), FSL (Jenkinson et al., 2012), AFNI (Cox, 1996), and ANTs (Avants et al., 2009). Within each run, functional volumes underwent slice timing correction and motion correction via rigid-body registration to the first volume. Then functional volumes were co-registered to the corresponding anatomical scan and spatially normalized to a standard rat brain template (Valdés-Hernández et al., 2011) using the symmetric diffeomorphic registration implemented in ANTs (Avants et al., 2009). The first 10 volumes were discarded. For the remaining volumes, we regressed out 2nd-order polynomial trends to remove slow scanner drift. Finally, the data were spatially smoothed using a 3D Gaussian kernel with a full width at half maximum (FWHM) of 1 mm.

### Seed-Based Correlation

We evaluated the seed-based functional connectivity using a General Linear Model (GLM). The seed was chosen from a list of predefined regions of interest (ROIs) along the established interoceptive pathway: including the dorsal vagal complex (DVC), thalamus (Thal), hypothalamus (Hypo), amygdala (Amy), insular cortex (IC), and cingulate cortex (Cg) from both the left and right hemispheres. As such, there were a total of 12 seed ROIs, as shown in (Supplementary Figure S1). These ROIs were defined using a standard MRI atlas of the rat brain (Valdés-Herńandez et al., 2011), except the DVC which was not accurately defined in the atlas. Thus, we manually delineated the DVC based on the Paxinos and Watson atlas (Paxinos & Watson, 2013) and the functional localization of the ascending vagal projection to the brainstem using neuronal tract tracing with manganese enhanced MRI (Oleson et al., 2023).

For each ROI, the BOLD signal was averaged across all voxels within the ROI to generate a seed time series. To estimate functional connectivity while rigorously controlling for motion artifacts, we constructed a GLM for every voxel in the brain. The model included the seed time course as the regressor of interest, alongside the six rigid-body motion parameters (translation and rotation) derived from the motion correction step as nuisance regressors. Prior to regression, both the seed and voxel time series were z-scored (i.e., demeaned and variance-normalized). Consequently, the resulting *β* coefficients represented the partial correlation between the seed and each voxel, independent of head motion.

Seed-based correlation maps for each ROI were entered into a second-level group analysis. Specifically, we performed a one-sample *t*-test (two-sided *p <* 0.05) on the voxel-wise partial correlation coefficients (*β* values) across all runs and animals within each group to identify voxels that exhibited significant functional connectivity.

### Conjunction Analysis

Subsequently, a conjunction analysis was performed to characterize the spatial convergence of functional connectivity across the different seed regions. For each seed, we generated a binary mask, assigning a value of 1 to voxels with significant functional connectivity and 0 otherwise. These masks were summed across all *N* seeds to generate a cumulative overlap map. The resulting overlap score at each voxel (ranging from 0 to *N* = 10) quantified the degree of functional convergence. Regions with high overlap scores defined the “core” regions that shared functional connectivity across seeds, distinguishing them from “peripheral” regions that exhibited correlations with specific seeds (but not all).

To assess the temporal stability of the conjunction analysis result, we performed a convergence analysis using truncated datasets. For each scan, we extracted the first 5, 10, 15, 20, or 25 minutes of the data. We mapped the interoception network using data from each truncated duration and quantified its Structural Similarity Index (SSIM) (Z. Wang et al., 2004) with the map based on data from 30 minutes.

## Results

### Functional Connectivity with DVC Extends from Brainstem to Cortex

Interoception relies on continuous signaling through peripheral and central neural pathways that interface at the dorsal vagal complex (DVC) in the lower brainstem (Powley, 2021). Within this complex, the nucleus of the solitary tract (NTS) receives afferent input relayed by vagal sensory neurons; the dorsal motor nucleus of the vagus (DMV) contains the motor neurons that transmit parasympathetic output; and the DMV receives direct input from NTS to enable closed-loop vago-vagal reflexive control (Travagli & Anselmi, 2016). As such, the DVC serves as an essential “gateway” for both the inflow of visceral sensation and the outflow of autonomic regulation. Given this pivotal role, we designated the DVC as the initial anchor for seed-based correlation analysis. We sought to determine whether functional connectivity of DVC is locally confined mainly to the vago-vagal reflex or extends to other subcortical and cortical regions for higher-order interoceptive processing.

Our analysis revealed that resting state functional connectivity with the DVC extended far beyond the brainstem, reaching higher-order regions along the established interoceptive hierarchy (Figure 1). These regions included the cerebellum, periaqueductal gray (PAG), specific nuclei in the thalamus and hypothalamus, as well as the insular, cingulate, and retrosplenial cortices, consistent with findings from prior neuronal tracing studies (Rinaman, 2010; Gasparini et al., 2020). In addition, DVC was also correlated with sensorimotor areas, including the primary visual, auditory, somatosensory, and motor cortices. Although the functional connectivity profiles of the left and right DVC were partially overlapped, we observed systematic distinctions. The left DVC was more strongly correlated with areas in the temporal and parietal association cortex, whereas the right DVC was more strongly correlated with subcortical and limbic areas, such as the thalamus, hypothalamus, hippocampus, and amygdala. Collectively, these findings demonstrate that the DVC anchors a distributed network of resting state functional connectivity, and this network is segregated into an “associative stream” linked to the left DVC and a “limbic stream” linked to the right DVC.

**Figure 1:**
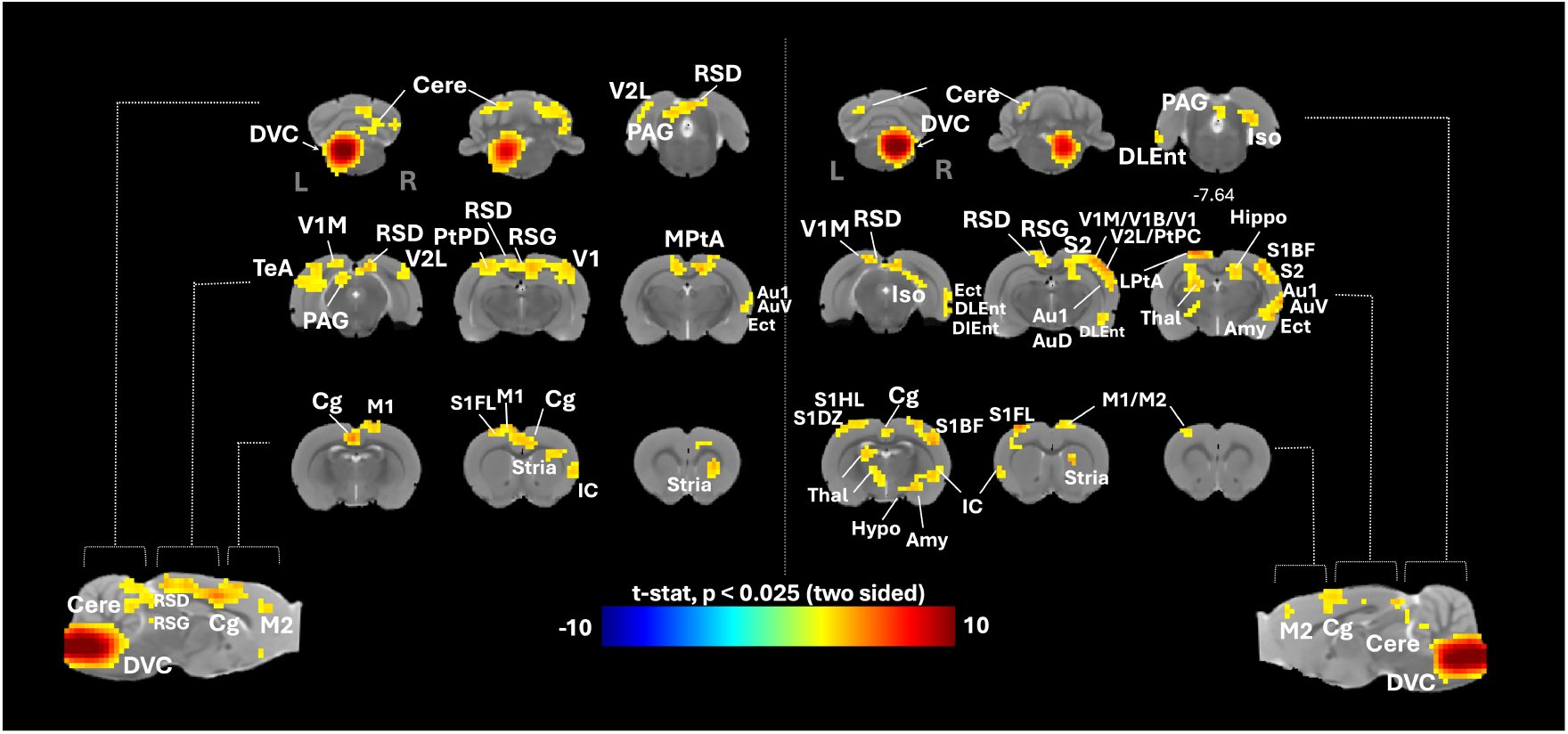
The dorsal vagal complex (DVC) was correlated with a distributed network. Based on data from *n* = 22 animals in the baseline cohort, the seed-based functional connectivity maps shown in the left and right panels correspond to the left and right DVC seeds, respectively. The color-coded maps display *t*-statistics from a one-sample *t*-test (two-sided, *p <* 0.025). The left on the image is also the left of the animal’s brain hereinafter. Annotations indicate major anatomical structures. See Supplementary Table S1 for abbreviations.

### The Interoception Network Exhibits Reciprocal Correlations

The functional connectivity of DVC allows for two distinct topological interpretations: a “common input” model versus a “cohesive network” model. In the former, the DVC acts as a central hub broadcasting common signals to target regions that otherwise function as isolated nodes. In the latter, these targets are not only correlated with the DVC but also mutually correlated with one another. To distinguish between these possibilities, we anchored the seed location to a set of core interoceptive regions, including the cingulate cortex, insular cortex, thalamus, hypothalamus, and amygdala, while separating each bilateral region into its left and right parts and using them as distinct seeds to fully capture the network topology. We generated a functional connectivity map for every seed and assessed the spatial overlap across different seeds.

The results provided strong support for the “cohesive network” model. Regardless of which region was chosen as the seed, the resulting functional connectivity map converged onto a similar spatial pattern (Figure 2). This spatial convergence uncovers a “highly integrated network” linking all core interoceptive regions, wherein functional connectivity to one region entails functional connectivity to all. For instance, seeds anchored in the thalamus or hypothalamus were robustly correlated with the insular and cingulate cortices, and vice versa. Such mutual correlations confirm that these regions are not merely parallel targets of a common source (the DVC in the lower brainstem), but collectively function as a cohesive network through dense, reciprocal interactions.

**Figure 2:**
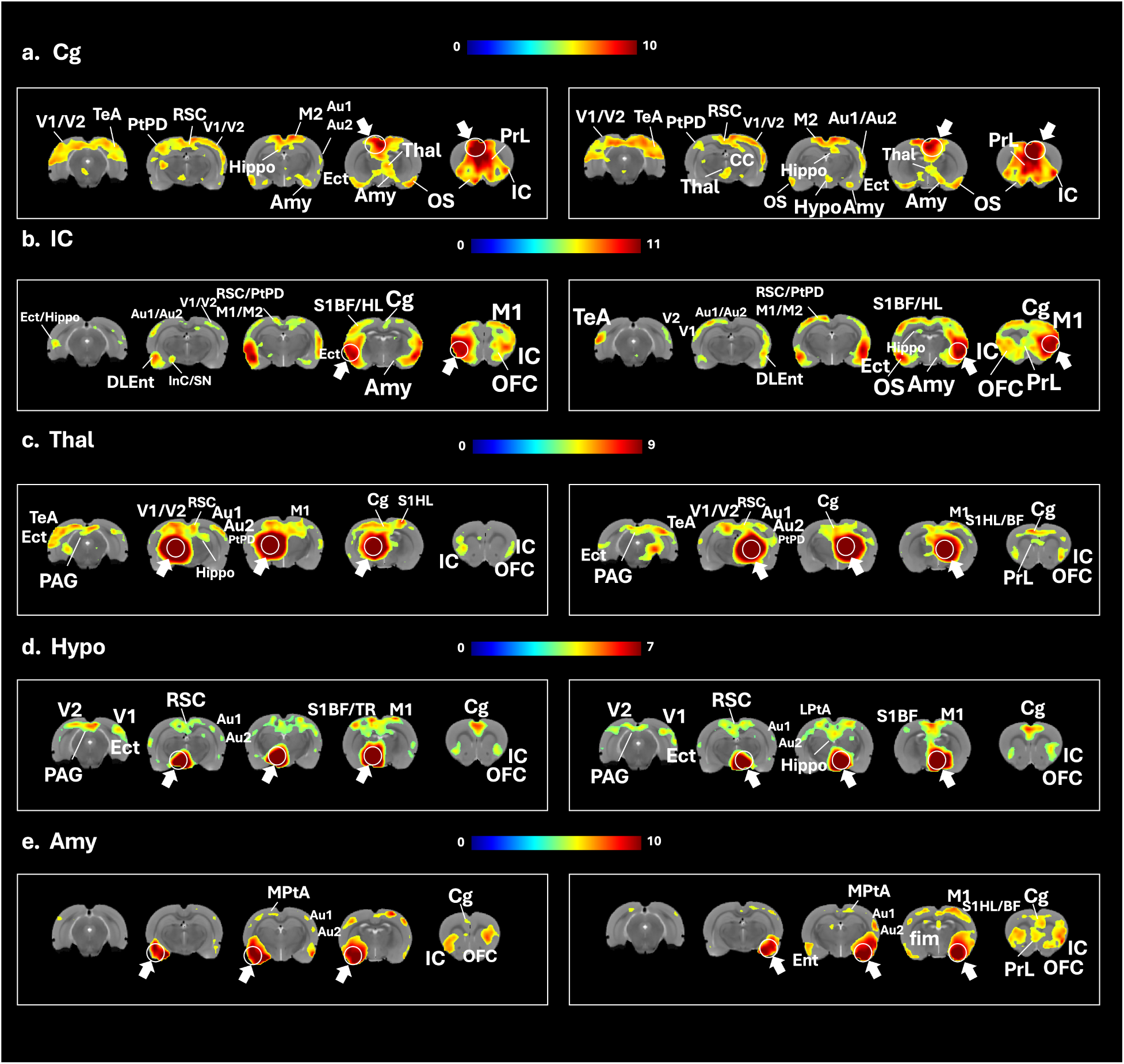
Reciprocal functional connectivity among core interoceptive regions. Seed-based correlation maps were generated for five additional key nodes (each split into the left and right parts) of the interoceptive pathway: (a) Cingulate cortex (Cg), (b) Insular cortex (IC), (c) Thalamus (Thal), (d) Hypothalamus (Hypo), and (e) Amygdala (Amy). White circles/arrows indicate the seed location for each panel. Regardless of the seed selection, the resulting functional connectivity maps converge onto a similar spatial pattern, demonstrating that these regions form a cohesive, reciprocally connected network rather than acting as isolated targets of the brainstem. Color bars represent *t*-statistics (two-sided, all maps were thresholded at *p <* 0.001).

To quantify this network integration, we generated a “conjunction map” visualizing the spatial convergence of functional connectivity across the different seed regions (Figure 3). In this map, the “overlap score” of each voxel corresponded to the number of distinct seeds that were functionally correlated with that voxel. This analysis effectively acted as a topological filter. High overlap scores highlighted the invariant network component, characterized by dense and mutual functional connectivity among its constituent nodes, whereas low overlap scores indicated variable functional connectivity specific to individual seeds. We found that the functional connectivity maps converged robustly onto the core interoceptive regions, confirming their strong reciprocal interactions. Moreover, this convergence extended beyond the predefined seed regions to encompass the striatum and widespread sensorimotor regions. Altogether, these distributed regions maintain reciprocal interactions and form a cohesive, integrated network.

**Figure 3:**
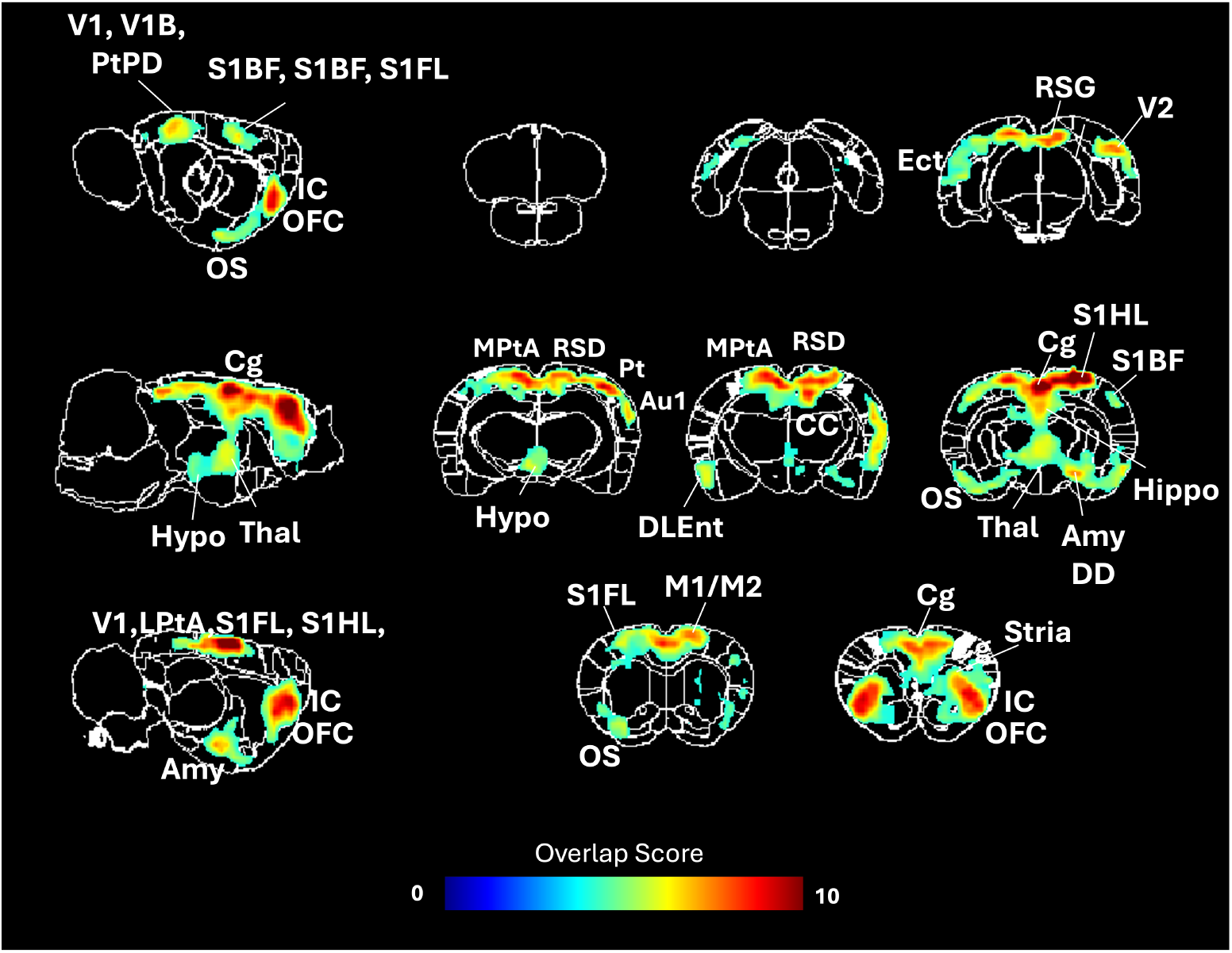
Topological convergence of the interoception network. A cumulative overlap map summarizes the spatial convergence of the 10 seed-based functional connectivity maps (bilateral seeds for Cg, IC, Thal, Hypo, Amy). The voxel intensity represents the Overlap Score (range: 0–10), defined as the number of seed regions that show significant functional connectivity with a given voxel. Regions with high overlap scores (red/yellow) represent network “hubs” that are consistently recruited by multiple regions along the interoceptive pathway, highlighting the network’s core with dense functional integration.

To assess temporal stability, we repeated this conjunction analysis with truncated versus full-length data. Results show that the interoception network revealed its pattern using data as short as 5 minutes. As more data were used, the pattern progressively converged at 15 minutes and beyond (Supplementary Figure S3). Thus, longer acquisitions primarily enhance the statistical stability rather than being compromised by autonomic shifts over time.

### The Interoception Network Depends on Feeding State

Herein, we refer to this network as the “interoception network” (Figure 3), given our hypothesis that it serves an interoceptive function. If this hypothesis is true, the functional topology of this network is expected to be sensitive to the varying physiological state. To test this, we compared animals in the fed and fasted conditions, corresponding to the digestive and inter-digestive phase of the gastrointestinal tract, respectively. We prioritized the gastric modality to capture distinct physiological states while avoiding direct manipulation of the cardiac or respiratory systems that might introduce vascular confounds to the fMRI signal. Specifically, we performed the conjunction analysis, computed voxel-wise overlap scores to measure functional integration, and compared these scores between fed and fasted conditions.

The results confirmed that the interoception network depends on the feeding state (Figure 4). In the fed condition, the network exhibited robust functional integration. Core interoceptive hubs and widespread sensorimotor cortices consistently showed high overlap scores, indicating that these regions maintained reciprocal functional connectivity with one another. In contrast, the degree of network integration was suppressed in the fasted condition. The spatial overlap in functional connectivity with individual nodes was overall weaker and more restricted. Relative to the fed state, the thalamus and sensorimotor cortices exhibited significantly weaker functional coupling, or even complete decoupling, with other regions in the network. An exception was the hypothalamus, showing stronger coupling within the rest of the network under the fasted condition. Quantitatively, the structural similarity of the network topology was higher between the fed and baseline cohorts (SSIM=0.7) than the similarity between the fasted and baseline cohorts (SSIM=0.63). As we previously reported using the same dataset (Cao et al., 2022), there were no significant differences in heart rate, respiratory rate, and SpO_2_ between the fasted and fed states, thereby ruling out potential vascular confounds from cardiac or respiratory fluctuations. These findings confirm the physiological relevance of the interoception network as shown in Figure 3, demonstrating its versatility to reconfigure its functional topology in response to shifts in digestive state and supporting our identification of this network as the “interoception” network.

**Figure 4:**
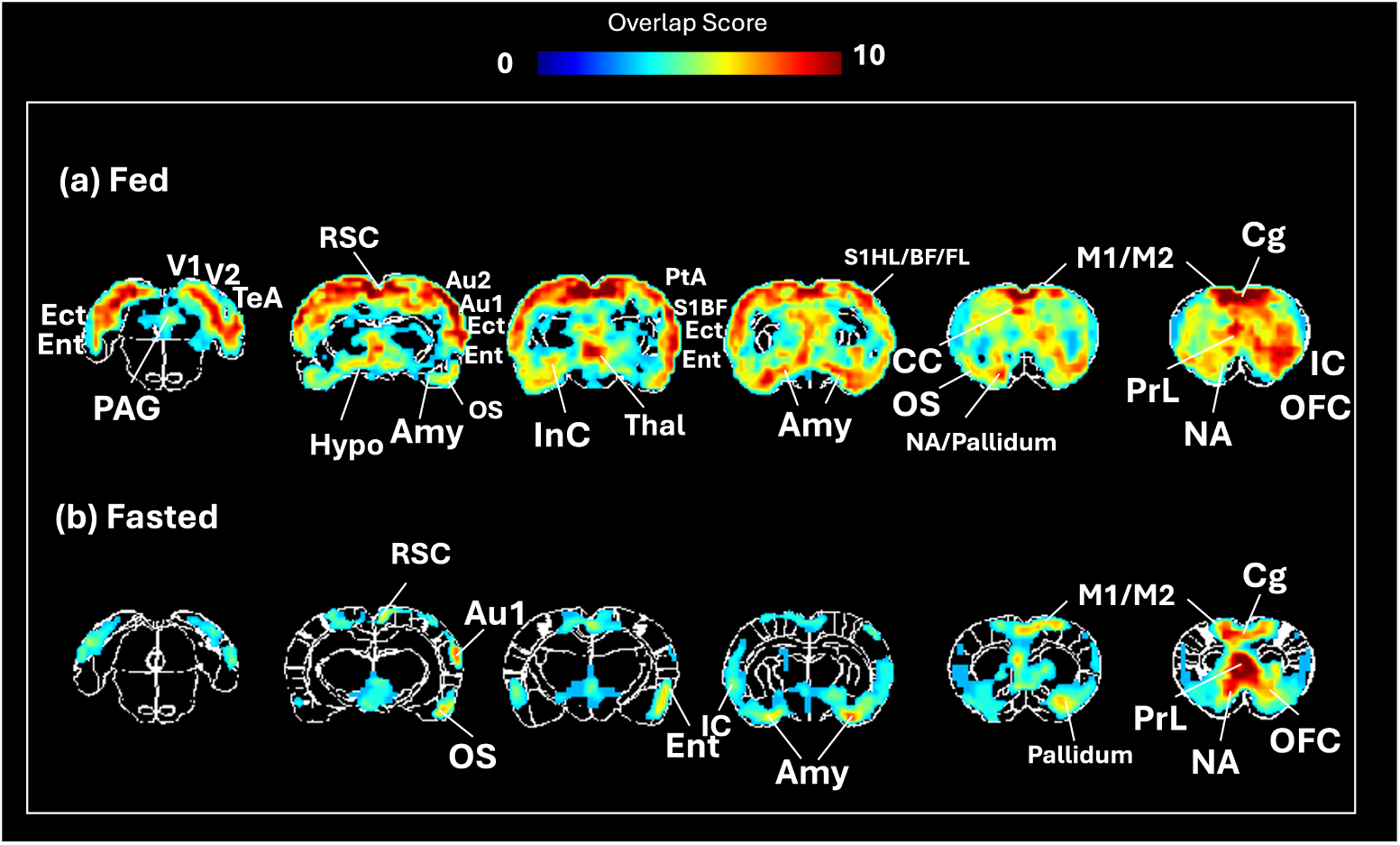
The interoception network was state-dependent. Conjunction analysis compared the network topology between the (a) fed (digestive phase) and (b) fasted (inter-digestive phase) conditions. The overlap scores (range: 0–10) indicate the degree of network integration.

### Bilateral Cervical Vagotomy Diminishes the Interoception Network

Our findings thus far establish that the interoception network is highly integrated via reciprocal functional connectivity and is responsive to changes in the feeding state. Next, we investigated whether the functional integration of this network relies on the vagus nerve, the primary neural pathway for rapid signaling between the brain and the stomach, among other organs.

To address this question, we surgically eliminated vagal signaling with bilateral cervical vagotomy. This procedure severed the vagal nerves between the brain and nearly all visceral organs, effectively blocking both vagal afferent input to the DVC and vagal efferent output from the DVC. We compared the functional topology of the interoception network in fed animals with either intact vagus nerves or cervical vagotomy to determine whether vagal integrity is a prerequisite for the network’s functional integration.

The results demonstrate that bilateral cervical vagotomy caused the interoception network to disintegrate (Figure 5). In contrast to the distributed and integrated topology observed in animals with intact vagus nerves, the network became severely fragmented in the absence of vagal signaling. This disintegration was characterized by two key features: 1) a complete disengagement of the sensory cortices from the rest of the network; and 2) a significant reduction in functional connectivity among the core interoceptive hubs. Although these hubs remained in the network, their coupling was weaker and less reciprocal, suggesting a profound reduction of functional integration. Quantitatively, the structural similarity of the network topology was higher between the non-vagotomized and baseline cohorts (SSIM=0.7) than between the vagotomy and baseline cohorts (SSIM=0.38).

**Figure 5:**
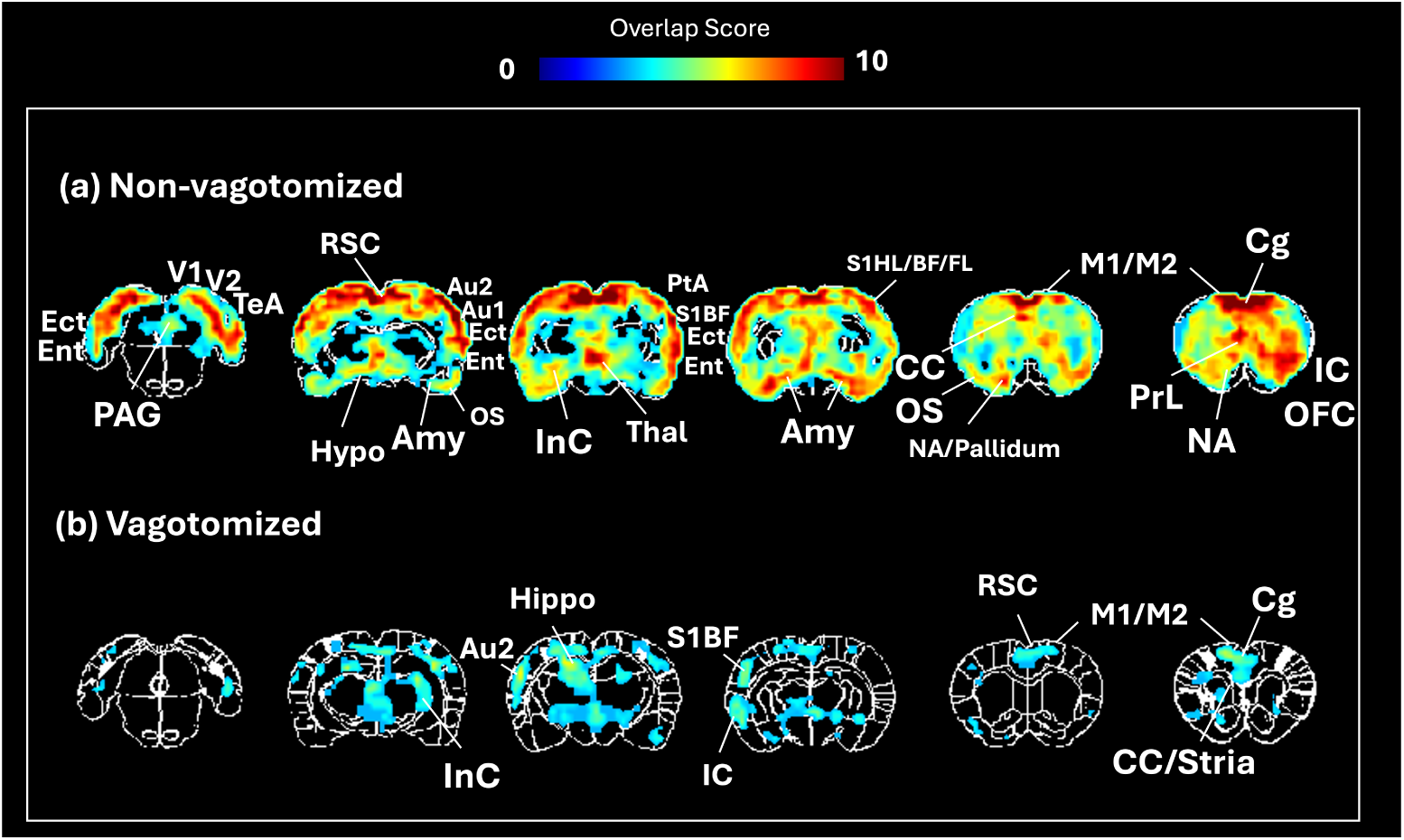
Vagal integrity was necessary for the functional integration of the interoception network. Comparison of network topology between (a) non-vagotomized (intact) and (b) vagotomized animals in the fed state. The overlap scores (range: 0–10) indicate the degree of network integration.

## Discussion

Using resting state fMRI in anesthetized rats, this study identifies a central interoception network (Figure 3), including dorsal vagal complex, parabrachial nucleus, hypothalamus, thalamus, amygdala, insula, and cingulate cortex, which is generally consistent with recent findings from human resting state fMRI studies (Kleckner et al., 2017; Zhang et al., 2025). Uniquely, we demonstrate that the interoception network is characterized by reciprocal functional connectivity among its constituent nodes. This “fully-connected” topology is not intrinsic to the brain but dependent on bodily states and peripheral neural signaling: it emerges robustly in the fed state but diminishes following fasting (Figure 4) or cervical vagotomy (Figure 5). In the following sections, we discuss the conceptualization of this network as an “integrative workspace”, the physiological interpretation of its state-dependence, the role of vagal signaling in maintaining brain-body interactions, and the generalization to cardiac and respiratory systems.

### Integrative Workspace for Brain-Body Interactions

The findings from this study lead us to posit that resting state networks are not merely intrinsic central networks, but dynamical systems coupled with the body’s physiological state. Central to this coupling is the interoception network (Figure 3), which maintains reciprocal correlations among its constituent nodes, including the dorsal vagal complex, parabrachial nucleus, hypothalamus, thalamus, amygdala, insula, cingulate cortex, as well as parts of the sensorimotor cortices. Given this “fully-connected” topology, engaging one node engages the entire network.

Therefore, we conceptualize the interoception network as an “integrative workspace” for brain-body interactions. This network can receive an incoming signal via any node, process the signal by engaging all nodes, and send an outgoing signal via any node. For instance, an ascending visceral signal entering this network via the DVC quickly recruits the entire network to generate an integrated representation of the physiological state, which is then broadcast to downstream systems to shape perception, emotion, cognition, and behavior. Conversely, the workspace is equally accessible for top-down regulation. Cognitive or emotional states entering the network via cortical nodes can recruit the entire network to synthesize descending regulatory signals that are broadcast to the viscera. In this framework, the interoception network acts as a flexible interface to balance physiological and mental states.

The interoception network is well positioned to interact with diverse neural and cognitive systems (Liu et al., 2023). Its nodes are widely recognized as “connector hubs” shared across different functional networks (van den Heuvel & Sporns, 2011; Power et al., 2013). For instance, the anterior insula and cingulate cortex are shared with the salience network (Seeley et al., 2007; Seeley, 2019), facilitating the balance of attention between internal and external states (Menon & Uddin, 2010). The amygdala and striatum are shared with the limbic system, associating interoception with emotion and motivation (Critchley & Garfinkel, 2017). The dorsal vagal complex and hypothalamus are shared with the autonomic control network to maintain homeostasis (Benarroch, 1993; Saper, 2002; Beissner et al., 2013). The interoception network also partially overlaps with the default-mode network (Lu et al., 2012), primarily at the retrosplenial cortex and medial prefrontal cortex. Quantitatively, about 36% of the voxels in the interoception network are also within the default-mode network (Supplementary Figure S2). By overlapping with other functional networks, the interoception network ensures that bodily states are embedded into the broader functional landscape of the brain.

While being highly integrative, the interoception network has dual streams: the left DVC anchored the “associative stream” (temporal and parietal association cortices); the right DVC anchored the “limbic stream” (amygdala and insula via subcortical nuclei), as shown in Figure 1. This lateralization is consistent with the distinct functional roles of the left and right vagus nerves. The right vagus is preferentially involved in regulating heart rate, while a recent study demonstrates that elevated heart rate drives affective behavioral state (e.g., anxiety) (Hsueh et al., 2023). Recent work has shown that the right vagus preferentially recruits the dopaminergic reward system for gut-induced motivation (Han et al., 2018). In contrast, the left vagus drives ascending neuromodulatory effects (Hulsey et al., 2017) to contextualize sensorimotor behavior and learning (Engineer et al., 2011).

Many resting state networks are preserved across species, including rodents, non-human primates, and humans (Pagani et al., 2023). Although rigorous counterparts for the interoception network requires further investigation, the network identified here in the rat brain is qualitatively similar to related seed-based correlational patterns observed in human resting-state fMRI (Kleckner et al., 2017; Zhang et al., 2025). In general, fMRI studies in animal models allow for invasive physiological perturbations, offering mechanistic insights that cannot be readily achieved in human fMRI studies.

### Dependence on the Physiological State

Given this notion of “integrative workspace”, the functional topology of the interoception network is not a static, intrinsic feature of the brain, but an emergent property associated with the physiological state. The robust integration observed in the fed condition demonstrates that the network is actively engaged in sensing and regulating gastrointestinal physiology. During the digestive phase, the stomach must coordinate complex motor events to facilitate gastric emptying (Goyal et al., 2019). The proximal stomach sustains tonic contractions to generate intragastric pressure; the distal stomach generates peristaltic contractions to grind and propel food toward the pylorus; and the pyloric sphincter must relax synchronously to permit transit into the duodenum (X. Wang et al., 2024; Di Natale et al., 2023). Simultaneously, the intestine absorbs nutrients and provides feedback to dynamically adjust the rate of gastric emptying (Maljaars et al., 2008). These coordinated events are not fully autonomous or solely managed by local circuits or the enteric nervous system (Di Natale et al., 2024), but rely on long-range gut-brain interactions that recruit the central interoception network into a cohesive, integrative system (Browning & Travagli, 2014; Powley, 2021).

In contrast, the inter-digestive (fasted) phase is relatively quiet. The stomach exhibits the migrating motor complex (Deloose et al., 2012), characterized by long periods of motor quiescence interspersed with brief periods of active contraction to clear out undigested residuals. Relative to the fed condition, the fasted condition imposes a significantly lower demand on the gut-brain neuroaxis, leading to a diminished level of integration within the central interoception network. However, an interesting exception is the hypothalamus, the brain’s “hunger center” (Morton et al., 2006), which maintained high functional connectivity even in the fasted state. Uniquely, the hypothalamus directly receives circulating hormones, e.g., ghrelin (Nakazato et al., 2001), thereby bypassing the neural pathway and maintaining independent access to the body’s metabolic state. This hormonal signaling pathway offers a plausible mechanistic explanation for our finding.

Our feeding paradigm is designed to be naturalistic, utilizing voluntary consumption rather than oral gavage. The pre-meal fasting ensures a consistent digestive state across animals, and the diet habituation conditions the animal for voluntary consumption. While these procedures are necessary for our experiments, we acknowledge that this feeding condition is not entirely equivalent to spontaneous eating. A potential concern is that our paradigm might be confounded by binge eating, which is primarily reward driven. However, we strictly cap the meal size to 5 g, precluding excessive consumption characteristic of binge eating. The dietary habituation further establishes a homeostatic need to drive normal eating behavior. Therefore, the observed difference between fed and fasted condition reflects naturalistic physiological processes, rather than transient psychiatric states.

### Active Vagal Signaling Sustains the Interoceptive Network

Our findings identify the vagus nerve as a primary pathway that sustains the functional integration of the interoception network. Surgical disruption of this pathway led to a profound reduction of the network’s reciprocal functional connectivity and a dissociation between sensory cortices and core interoceptive regions. Consistent to this finding, prior research also demonstrates in rats that vagotomy diminishes stomach-brain synchronization (Cao et al., 2022), which is observable in humans (Rebollo et al., 2018; Choe et al., 2021; Levakov et al., 2023). Together, these findings confirm that resting state functional connectivity is not solely an intrinsic property of the central nervous system but must be interpreted in the context of the peripheral nervous system, which supports continuous bidirectional signaling between the brain and visceral organs.

The reported effect of the vagotomy on the interoception network should be interpreted in the context of our experimental timeline and the physical properties of our test meal. Small liquid meals can trigger rapid, transient gut-brain neural signaling (Essner et al., 2025; Beutler et al., 2017; Kaelberer et al., 2018). However, our animals consumed a 5-g semi-solid meal. Under normal physiological conditions, such a meal requires at least four hours to fully empty from the stomach, and the use of anesthesia further prolongs the emptying time (Lu et al., 2017; X. Wang et al., 2026). Given our experimental timeline, the vagotomy was performed approximately 15 minutes after the meal. Its effects on the central interoception network reflect sustained vagally-mediated gut-brain interactions during the postprandial phase. This phase is not stationary but involves a physiological cascade, ranging from initial accommodation and food trituration to transpyloric transit, duodenogastric feedback, and distal nutrient sensing. It is likely through the vagus that the interoception network maintains the central monitoring and regulation of this multi-stage process, which lasts for hours unlike the initial, rapid sensory activation during or immediately after food intake (Essner et al., 2025). This is, however, speculative, awaiting future studies that combine data recorded from both the gut and the brain.

It is worth noting that the vagus nerve is a major, but not the exclusive, peripheral pathway for interoception. For example, the gut is also innervated by spinal nerves (Zhang et al., 2022), which provide afferent signaling through the dorsal root ganglia and sympathetic control via the celiac ganglion, among others. Even after the complete elimination of vagal signaling by bilateral cervical vagotomy, the brain and viscera maintain some degree of communication through these spinal pathways, which may mediate compensatory responses to visceral sensation (Sanvanson et al., 2019). However, the fact that the interoception network was significantly attenuated despite these remaining connections suggests that the vagus nerve plays an indispensable and likely major role in maintaining the “integrative workspace” topology.

Its dependence on vagal integrity establishes the interoception network as a potential biomarker to evaluate and guide vagus nerve stimulation (VNS) (Lu et al., 2018). If the “fully-connected” functional topology represents the neural signature of normal postprandial physiology, this specific network state can serve as a target biomarker for using VNS to normalize pathophysiology. Thus, we propose a “tune-and-match” strategy for optimizing VNS based on brain scans or recordings (Cao et al., 2017, 2019, 2021). The stimulation parameters (e.g., frequency, pulse width, amplitude) could be dynamically tuned in real-time until the brain network topology matches the normal, integrated “target” topology. This approach moves beyond any heuristic or “one-size-fits-all” stimulation protocols to potentially enable personalized, closed-loop therapies grounded in the objective evidence for engaging the central interoception network in restoring brain-body communication.

### Generalization to Cardiac and Respiratory Systems

Arguably, the interoception network serves as a generalized integrative workspace for multi-organ regulation, extending beyond the gut to include the cardiovascular and respiratory systems, etc. The network’s core nodes, specifically the DVC, insula, and anterior cingulate cortex, are established centers for cardio-respiratory control. However, validating this hypothesis via fMRI is non-trivial. Cardiac and respiratory fluctuations, such as changes in heart rate and respiratory volume per time (Birn et al., 2006; Shmueli et al., 2007), induce systemic hemodynamic shifts, such as carbon dioxide (CO_2_)-induced vasodilation (Wise et al., 2004). These fluctuations are often treated as “physiological noise” and removed during preprocessing (Birn et al., 2008; Chang et al., 2009; Kasper et al., 2017). It remains a critical challenge to disentangle whether any observed correlations reflect genuine neurogenic interoception or passive vasogenic artifacts (Bright et al., 2017; Liu, 2016; Liu et al., 2023), or effects of arousal (Bolt et al., 2025). Resolving this ambiguity awaits future studies utilizing strict experimental controls, e.g., clamping end-tidal CO_2_ (Golestani & Chen, 2020), or mechanistic modeling grounded in the theory of interoception, e.g., predictive coding (Barrett & Simmons, 2015; Choi et al., 2025).

### Limitations and Considerations

A methodological limitation of the current study is the absence of a sham-operated control group. Instead, the vagotomy cohort was compared against a control group of intact animals that did not undergo any surgical procedures. Consequently, we cannot definitively separate the specific physiological effects of severing the vagus nerve from the non-specific effects of surgical stress. While we cannot completely dismiss the impact of generalized surgical stress, the spatial distribution of the observed functional decoupling is more consistent with established vagal-brainstem-cortical pathways rather than a uniform, brain-wide suppression.

Anesthesia alters autonomic physiology. In this study, fMRI was performed in anesthetized rats, while it is not practical to perform vagotomy in awake rats. To mitigate the confounding effects of anesthesia, animals were maintained in a stable physiological condition, ensured by continuous monitoring of SpO_2_, body temperature, heart rate, and respiration. Across the fed, fasted, and vagotomized conditions, we did not observe significant differences in heart rate, respiration rate, or the global fMRI signal, although the heart rate variability was lower in the vagotomized and fasted conditions relative to the fed condition (Cao et al., 2022). Furthermore, it remains challenging to definitively pinpoint the specific physiological origins of the observed network alterations. The changes induced by systemic vagotomy likely reflect a complex mixture of cardiac, respiratory, and gastrointestinal effects, as well as intertwined vascular and neuronal responses. Disentangling these overlapping mechanisms will require future studies that combine fMRI with simultaneous, direct measurements of neuronal and vascular activity.

While we focus on interoception, the effect of vagotomy may not be exclusively confined to the interoception network. The interoception network overlaps with several other functional networks (Liu et al., 2023), such as the salience network. The effects upon the interoception network can extend further to other networks through their overlapped regions. While it is outside the scope of this paper, a comprehensive, whole-brain characterization of all resting state networks is expected to be valuable for understanding the general role of vagal signaling in spontaneous brain activity.

In this retrospective study, we pooled female Fischer and male SD rats to enhance the statistical power for mapping the baseline interoception network, while the core topologies did not significantly differ between them. All physiological interventions were evaluated in separate cohorts of only male SD rats, ensuring the observed effects of the feeding state and the bilateral vagotomy are not confounded by sex or strain. However, this study was not designed or statistically powered to properly test the effects of sex or strain as a biological variable, future prospective studies are required to investigate potential sex- and strain-specific differences in the interoception network.

## Conclusion

In conclusion, resting state networks are not merely intrinsic features of the brain, but dynamical systems coupled to the body’s internal organs. We identify the interoception network as a specialized functional system for this coupling, characterized by a “fully-connected” topology with reciprocal functional connectivity among the dorsal vagal complex, parabrachial nucleus, hypothalamus, thalamus, amygdala, insula, and cingulate cortex. This architecture functions as an “integrative workspace”, linking interoception with broader neural processes. Crucially, this network is actively sustained by vagal signaling and is highly sensitive to the digestive state. Our findings redefine the so-called resting state as an embodied phenomenon, establishing the interoception network as a critical, state-dependent interface between the brain and the body.

## Data and Code Availability

Data and code will be available via https://doi.org/10.7302/jwdg-4w60.

## Author Contributions

FA conducted data analysis. XW contributed to data acquisition. ZL conceived the study. FA and ZL wrote the paper. FA, XW, and ZL reviewed and revised the paper.

## Acknowledgment

This study has been supported by grants from the National Institutes of Health: OD023847, OD030538, and AT011665. The authors appreciate the valuable discussions with Dr. Ulrich Scheven and would like to acknowledge Dr. Jiayue (Cherry) Cao for acquiring the datasets and for her scientific insight which continues to inspire this work.

## Supplementary Materials

**Figure S1:**
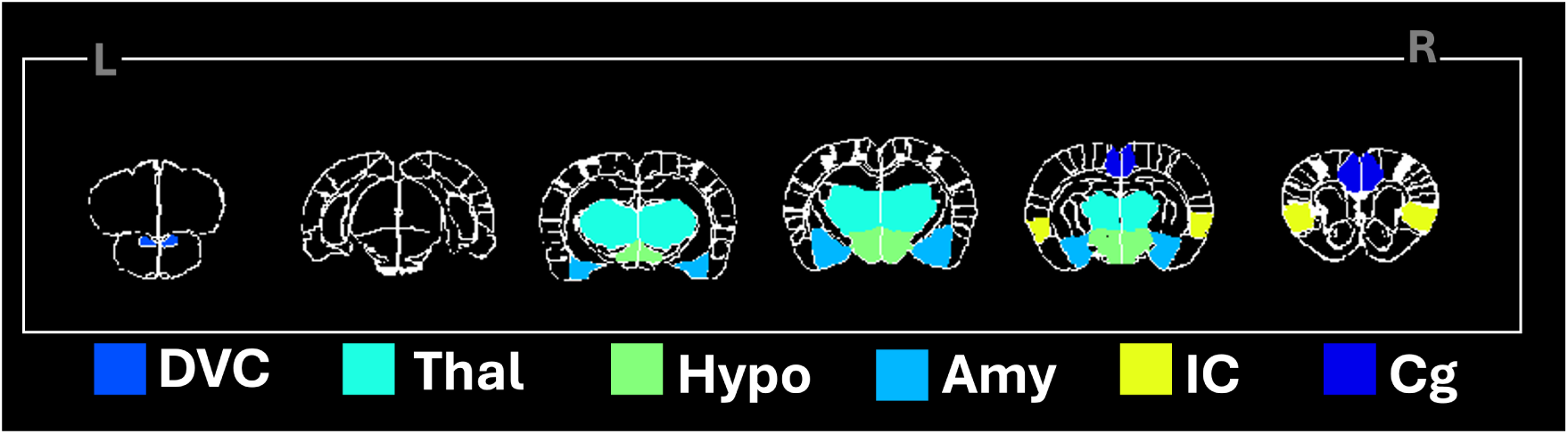
Anatomical ROIs are overlaid on representative brain slices: dorsal vagal complex (DVC), thalamus (Thal), hypothalamus (Hypo), amygdala (Amy), insular cortex (IC), and cingulate cortex (Cg)

**Figure S2:**
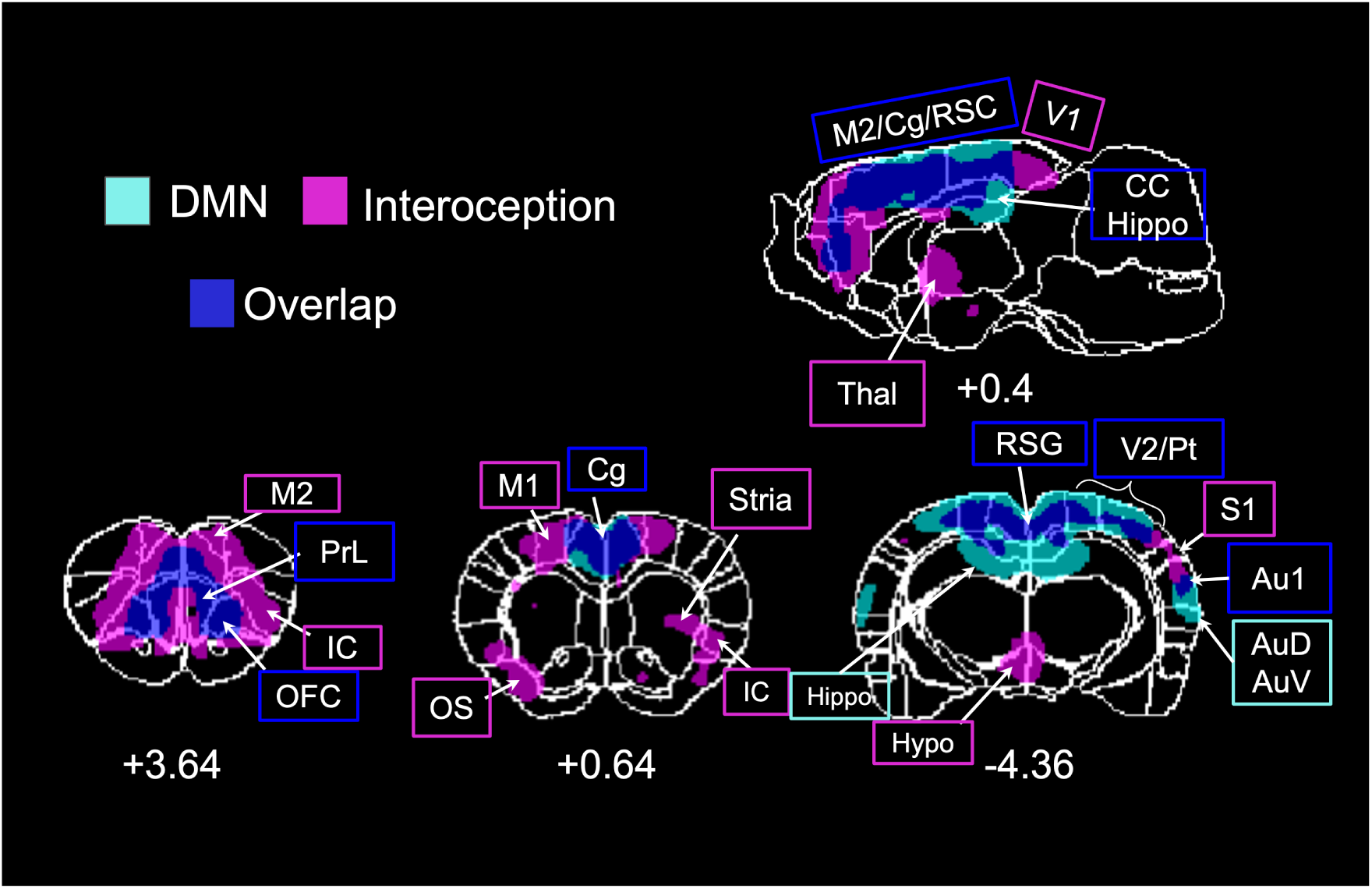
Partial overlap between the interoception network and the default mode network (DMN) in the rat brain. Cyan denotes regions assigned to the DMN, pink denotes regions assigned to the interoception network, and blue indicates overlapping voxels. In total, 35.5% of voxels in the interoception network are also part of the DMN. The DMN mask is derived from the DMN published in (Lu et al., 2012).

**Figure S3:**
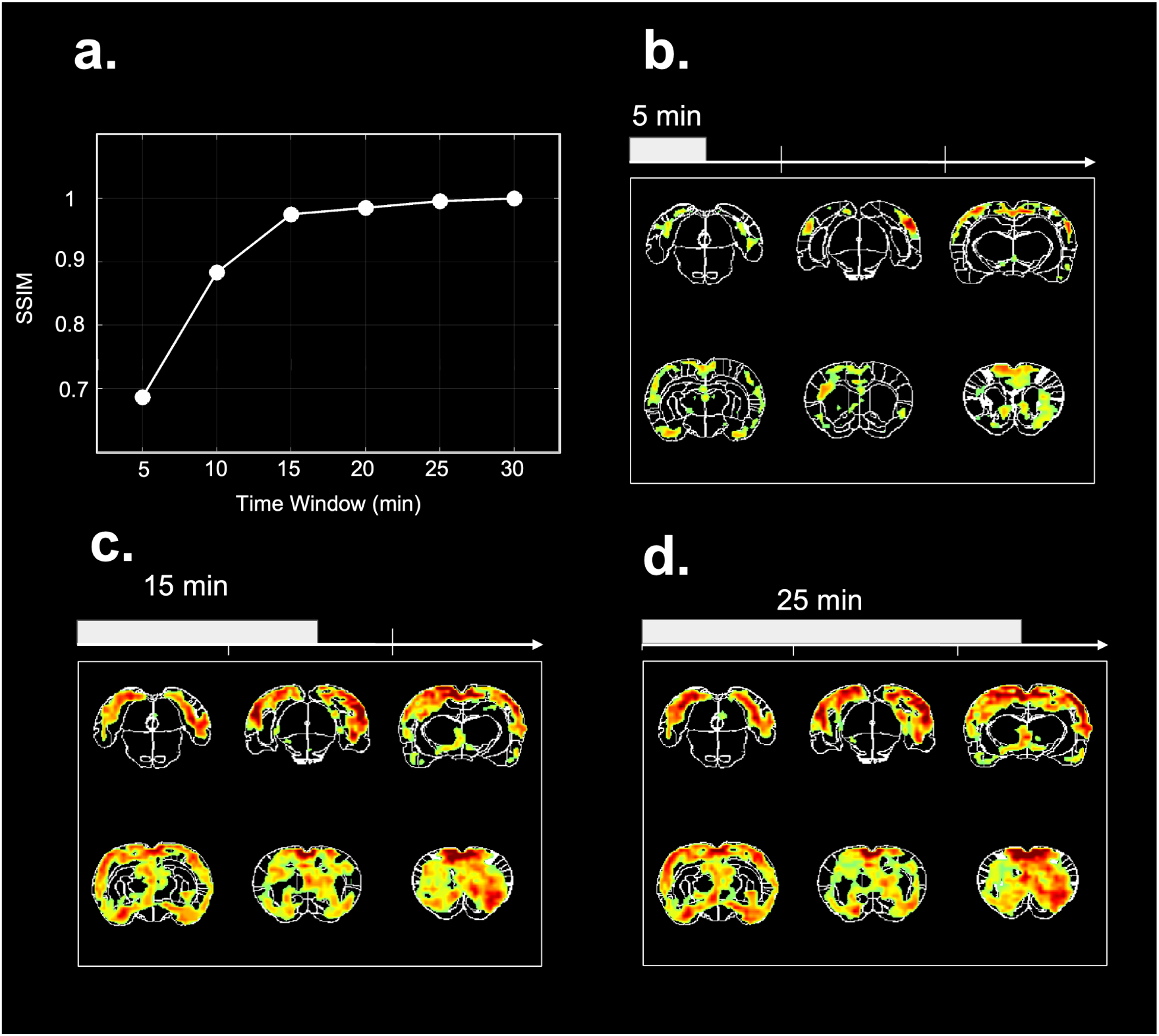
a.Structural Similarity Index (SSIM) at each time window relative to the 30 min window. b, c and d are conjunction maps computed with 5, 15, and 25 min windows respectively

**Table S1:** Brain region abbreviations and anatomical definitions used in this study.

| Abbreviation | Full region name |
| --- | --- |
| Amy | Amygdala |
| Au1 | Primary auditory cortex |
| AuV | Secondary auditory cortex, ventral area |
| Cg | Cingulate cortex |
| Cere | Cerebellum |
| DD | Diagonal domain |
| DIEnt | Dorsal intermediate entorhinal cortex |
| DLEnt | Dorsolateral entorhinal cortex |
| DVC | Dorsal vagal complex |
| Ect | Ectorhinal cortex |
| fim | Fimbria |
| Hippo | Hippocampus |
| Hypo | Hypothalamus |
| IC | Insular cortex |
| InC | Internal capsule |
| Iso | Isocortex |
| LPtA | Lateral parietal association cortex |
| M1 | Primary motor cortex |
| M2 | Secondary motor cortex |
| MPtA | Medial parietal association cortex |
| OFC | Orbitofrontal cortex |
| OS | Olfactory structures |
| PAG | Periaqueductal gray |
| PrL | Prelimbic area |
| PtPC | Parietal cortex, posterior area, caudal part |
| PtPD | Parietal cortex, posterior area, dorsal part |
| RSD | Retrosplenial dysgranular cortex |
| RSG | Retrosplenial granular cortex |
| S1BF | Primary somatosensory cortex, barrel field |
| S1DZ | Primary somatosensory cortex, dysgranular zone |
| S1FL | Primary somatosensory cortex, forelimb region |
| S1HL | Primary somatosensory cortex, hindlimb region |
| SN | Substantia nigra |
| Stria | Striatum |
| TeA | Temporal association cortex |
| Thal | Thalamus |
| V1 | Primary visual cortex |
| V1B | Primary visual cortex, binocular area |
| V1M | Primary visual cortex, monocular area |
| V2L | Secondary visual cortex, lateral area |

